# Comprehensive analysis of mutational signatures in pediatric cancers

**DOI:** 10.1101/2021.09.28.462210

**Authors:** Venu Thatikonda, S. M. Ashiqul Islam, Barbara C. Jones, Susanne N. Gröbner, Gregor Warsow, Barbara Hutter, Daniel Huebschmann, Stefan Fröhling, Mirjam Blattner-Johnson, David T.W. Jones, Ludmil B. Alexandrov, Stefan M. Pfister, Natalie Jäger

**Author notes:** **Corresponding authors** Correspondence to Stefan Pfister and Natalie Jäger.

## Abstract

Analysis of mutational signatures can reveal the underlying molecular mechanisms of the processes that have imprinted the somatic mutations found in a cancer genome. Here, we present a pan-cancer mutational signatures analysis of single base substitutions (SBS) and small insertion and deletions (ID) in pediatric cancers encompassing 537 whole genome sequenced tumors from 20 molecularly defined cancer subtypes. We identified only a small number of mutational signatures active in pediatric cancers when compared to the previously analyzed adult cancers. Further, we report a significant difference in the proportion of pediatric tumors which show homologous recombination repair defect signature SBS3 compared to prior analyses. Correlating genomic alterations with signature activities, we identified an association of *TP53* mutation status with substitution signatures SBS2, SBS8, SBS13 and indel signatures ID2 and ID9, as well as chromothripsis associated with SBS8, SBS40 and ID9. This analysis provides a systematic overview of COSMIC v.3 SBS and ID mutational signatures active across pediatric cancers, which is highly relevant for understanding tumor biology as well as enabling future research in defining biomarkers of treatment response.

## Main

Childhood cancers have a lower incidence rate when compared to the overall incidence of adult cancers. Nevertheless, cancer remains one of the leading causes of death by disease amongst children^1^. Growing research evidence suggests that childhood cancers are significantly different in terms of molecular features and therapy response when compared to their adult counterparts. Most prominently, recent pan-cancer analyses revealed that pediatric cancers show a significantly lower mutation burden in contrast to common adult cancers^2,3^. However, the knowledge of underlying mutational processes that contribute to the somatic mutation burden and tumor development in pediatric cancer is still limited.

Somatic mutations including single base substitutions, small insertions and deletions (indels), copy number changes, and other genomic rearrangements can be caused solely by endogenous processes (*e*.*g*., defects in DNA repair, errors in DNA replication, damage due to reactive oxygen species, *etc*.), or by the additional influence of exogenous causes such as exposure to ultra-violet light, tobacco smoking, and numerous others^4^. Previous pan-cancer analyses mainly focused on adult cancer types, employed mathematical models based on nonnegative matrix factorization (NMF) to identify patterns of somatic mutations, termed mutational signatures and utilized these patterns to infer underlying mutational processes^5^. Since then, mutational signatures have been used to understand tumor development, to identify gene alterations associated with mutational processes and importantly, as biomarkers for predicting treatment response^6,7,8,9,10,11,12^. A total of 45 single base substitution signatures (SBS) and 17 small insertion/deletion (ID) signatures were identified in a recent analysis as part of the pan-cancer analysis of whole genomes (PCAWG) consortium^13,14^, updating and expanding the previous set of reference mutational signatures (*i*.*e*., mutational signatures in COSMIC v.2).

Previous studies of individual pediatric tumor types and pan-cancer approaches analyzed mutational signatures as part of molecular tumor landscape analyses using the COSMIC v.2 reference signatures^15,16,17,18,19,20,23^. Here, we carried out an extensive analysis encompassing signatures of single base substitution and, for the first time, signatures of small insertion/deletions in 537 whole genome sequenced pediatric tumor-normal pairs. These were subsequently compared with COSMIC v.3^14^ signatures to identify overlap with the latest set of known mutational signatures.

## Results

### Somatic mutation frequencies across twenty pediatric cancer types

A dataset of 537 whole genome sequenced tumor-normal pairs (the PedPanCan (PPC-WGS) cohort, https://www.kitz-heidelberg.de/en/research/datacommons/pedpancan/) was compiled from a previously published study^2^, spanning 20 molecularly defined entities of childhood cancer (Fig. 1a). Single base substitutions (SBS) and small insertions and deletions (IDs) were identified using an updated in-house mutation calling pipeline as compared with the previous analysis, with minor variations mostly in the calling strategy of indels (Supp. Fig. 1a). In total, across the cohort we identified 2,712,521 SBS and 270,928 IDs. The number of SBS mutations per megabase (median: 0.24; range: 0.0035 - 645.70) and IDs (median: 0.033; range: 0.00142-46.64) was highly variable both across individual tumors as well as across different cancer types (Fig. 1a, Supp. Table 1). The lowest overall somatic mutation burden was observed in pilocytic astrocytoma (median: 0.0429 mutations per megabase) and the highest in osteosarcoma (median: 1.17). Although the indel mutation burden per tumor was low, numbers of SBS and IDs were significantly correlated across tumors (Fig. 1b). As previously described in different childhood and adulthood cancer types, the mutation burden of both SBS and IDs is also clearly correlated with age in this pediatric cohort (Supp. Fig. 1b). A small number (n=3) of tumors classified as high-grade glioma with germline alterations in DNA mismatch repair genes (*MSH6, PMS2*) showed a hyper-mutator phenotype^2^, with 78.98 SBS and 2.63 ID mutations/Mb, respectively (Fig. 1b).

**Figure 1:**
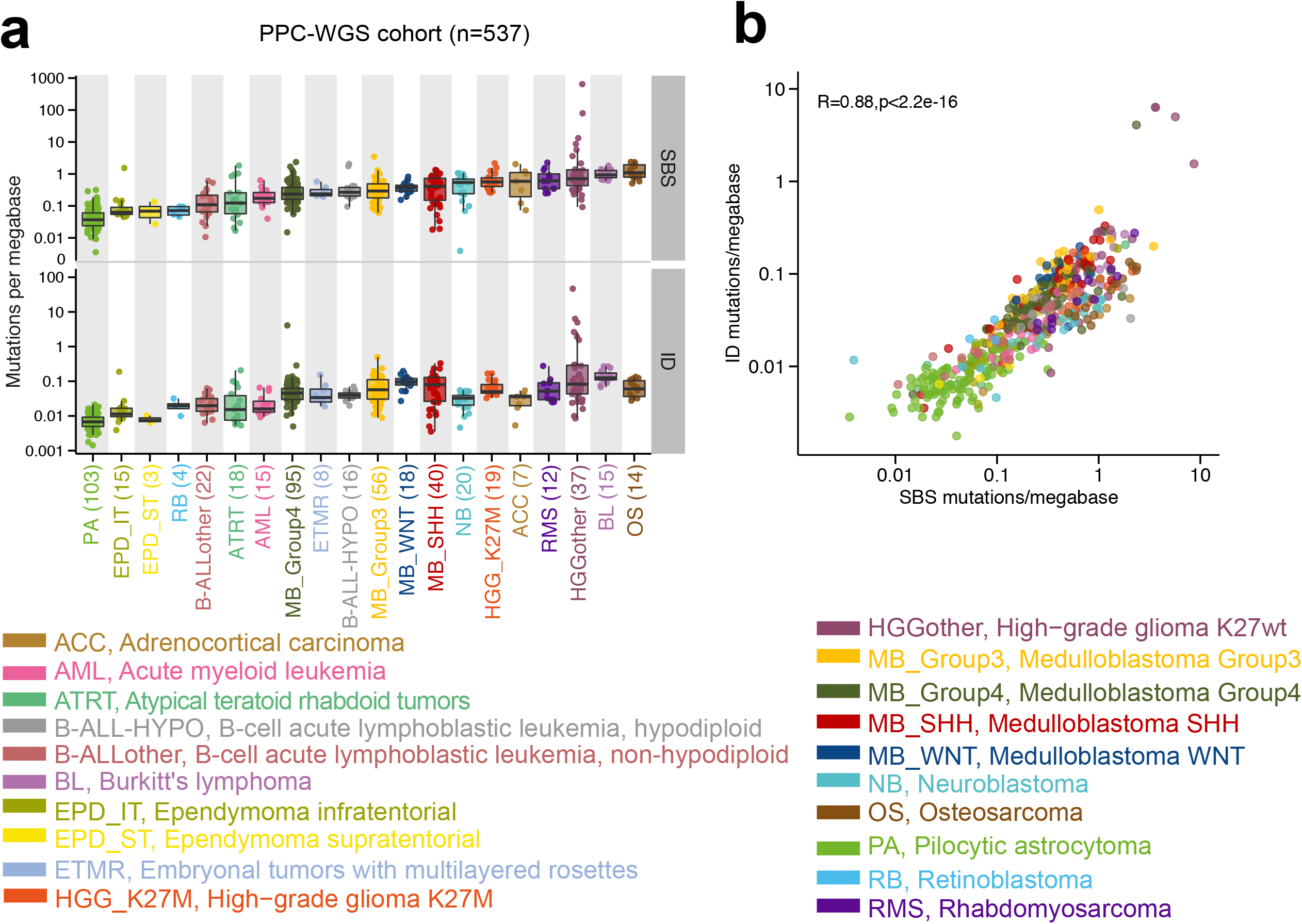
Mutation burden of SBSs and small indels (IDs) across 20 pediatric tumor types. a) Single base substitution (SBS) and small insertion/deletion (ID) mutation burden of the pediatric pan-cancer whole-genome sequenced (PPC-WGS) cohort. The numbers of samples for each tumor type are shown next to the labels. Each dot represents one tumor sample. Tumor types are ordered by the median numbers of single-base substitutions. b) Correlation between SBS and ID mutations per megabase across the cohort.

### SBS and ID signature activities

To extract mutational signatures active in this pediatric pan-cancer cohort, we generated 96-context SBS and 83-context ID mutational catalogues (Fig. 2a) and utilized an approach based on nonnegative matrix factorization, as previously described^5,21,14^. Due to the very low number of double base substitutions per pediatric tumor (data not shown), we only extracted signatures from SBS and IDs.

**Figure 2:**
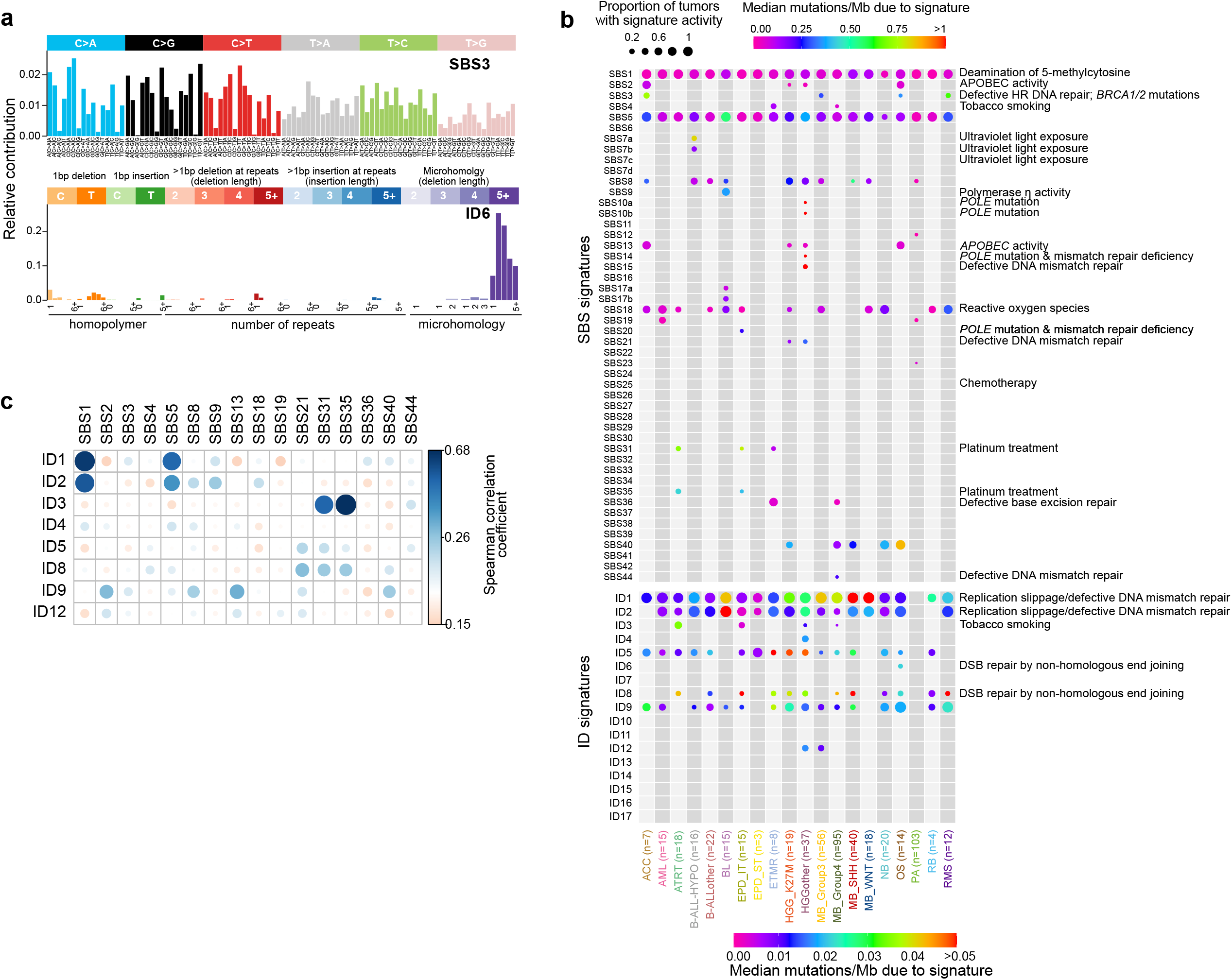
SBS and ID signature activity across pediatric cancers. a) Example profiles of SBS and ID signatures extracted from the PPC-WGS cohort that are similar to COSMIC v.3 SBS3 and ID6 signatures. b) The number of mutations contributed by each mutational signature to the PPC-WGS tumors. Circle size indicates fraction of tumors with signature activity in a cancer type and the color indicates the median number of mutations per megabase due to a signature in a specific entity. c) Correlation between SBS and ID signature activities.

In total, 27 SBS signatures were extracted that matched the COSMIC v.3 SBS signatures with a minimum cosine similarity of 0.9 (Fig. 2b, Supp. Fig. 2a). Amongst these, SBS1 and SBS5 were present in 97.5% and 96.1% of samples across the cohort, respectively (Fig. 2b, Supp. Table 2, Supp. Table 3). As described in adult cancers and a small fraction of pediatric brain tumors, the clock-like nature of SBS1 and SBS5 was also observed in this cohort, with a significant correlation of signature activity with age at diagnosis^7^ (Supp. Fig. 3a). An additional signature with unknown etiology, namely SBS40, is similar to SBS5 (cosine similarity =0.83)^14^. SBS40 was found to be active in five pediatric cancer types and 20% of tumors across the cohort (Fig. 2b, Supp. Table 3) and was also correlated with age at diagnosis (Supp. Fig. 3a), suggesting it may be an additional clock-like signature.

SBS2 and SBS13 were reported to be mainly due to the activity of APOBEC enzymes^14^ and in this cohort they were found to be present only in adrenocortical carcinoma (ACC), high-grade glioma and osteosarcoma (Fig. 2b, Supp. Table 2). In a cross-cohort comparison, mutations attributed to SBS2 and SBS13 were significantly higher in *TP53* germline mutated tumors compared to tumors with somatic or no *TP53* mutations (Supp. Fig. 3b). This observation is in line with previous studies which identified a link between p53 loss and elevated *APOBEC3B* expression^22,23^. However, not all *TP53* germline mutated tumors in this cohort showed SBS2 or SBS13 activity, including the *TP53*-defined subtype of SHH-medulloblastoma (Supp. Table 2 & Table 13).

Ultraviolet light (UV) exposure signatures SBS7a and SBS7b^14^ were identified in tumors of hypodiploid B-cell acute lymphoblastic leukemia (B-ALL-HYPO) (Fig. 2b), but not in non-aneuploid tumors. UV light affected tumors show an enrichment of dipyrimidine substitution mutations (CC>TT). Consistently, SBS7a and SBS7b positive tumors show a significantly higher number of CC>TT doublet base substitutions when compared to SBS7a and SBS7b negative tumors in the B-ALL-HYPO subtype (Supp. Fig. 3c). Although multiple independent studies have recently identified signatures apparently linked with UV light exposure in pediatric B-ALL^3,45^, the exact mechanism how it contributes to leukemogenesis, and whether UV light is really the cause or rather the signature is mimicked by another process, remains to be elucidated. In 1.5% of tumors (8/537; ETMR and Group4 medulloblastoma) we identified signature SBS4 (Fig. 2b, Supp. Table 2), which is proposed to be the result of DNA damage caused by tobacco smoking^14^. To the best of our knowledge, these tumors were derived from untreated, primary pediatric tumors and we hypothesize that the C>A mutation pattern consistent with SBS4 may potentially be due to a different exogenous mutational process generating a similar signature of mutations as tobacco smoking. No common germline or somatic mutations were identified in these eight brain tumors with SBS4 activity.

Other relatively frequent signatures across cancer types included SBS8 and SBS18 (Fig. 2b). Signature 8 from COSMIC v.2 was proposed to be due to homologous recombination repair pathway gene mutations^2^. We did not identify any such mutations in SBS8 positive pediatric tumors in this analysis. However, the number of mutations attributed to SBS8 was significantly higher in tumors with germline or somatic *TP53* mutations compared to wildtype tumors (Supp. Fig. 3d). Recent evidence suggests that the SBS8 signature is due to the DNA damage caused by late replication errors^24^. SBS18 is the result of DNA damage caused by reactive oxygen species^14^ and is observed in 12 out of 20 cancer types analyzed in this cohort (Fig. 2b). Amongst these 12 cancer types, the highest fraction of tumors with SBS18 signature activity were observed in neuroblastoma (85%, NB) and rhabdomyosarcoma (83%, RMS) (Supp. Table 4). Further, we found high activity of SBS18 in MYCN-amplified tumors in a cross-cohort analysis (Supp. Fig. 3h). However, in a per-cancer-type analysis of NB, RMS and Group3 medulloblastoma, tumors in which MYCN amplification is common, we did not identify a significant SBS18 association with MYCN status.

Mutations attributed to signature SBS9 are due to DNA damage caused by polymerase eta activity and were observed only in Burkitt’s lymphoma (BL) in this cohort. As previously described, SBS9 activity is found in BL with immunoglobulin gene hypermutations^36^ (Supp. Fig. 3e). Signature 10 from COSMIC v.2 has been split into SBS10a and SBS10b in the COSMIC v.3 signatures^14^. Mutations from high-grade glioma hyper-mutator tumors were attributed to both SBS10 signatures, as well as SBS14 and SBS15 (Fig. 2b). In Group4 medulloblastoma, we identified a novel SBS signature (termed Group4MB-SBS96D) with elevated T>A mutations in the context of C<u>T</u>T/T<u>T</u>T (Supp. Fig. 3f, Supp. Table 5). This signature contributed a large fraction of somatic point mutations to two tumors (3,623 in ICGC_MB174 and 1,458 in ICGC_MB175, respectively). However, we did not identify any molecular features common and specific to these tumors. Further investigation on the processing of tumor material revealed that the surgery for these two patients was performed on the same day in the same hospital, and therefore we suspect that the Group4MB-SBS96D signature is an additional artefact signature.

In addition, we identified SBS21 in high-grade glioma, which is the result of defective DNA mismatch repair, SBS36 in ETMR and Group4 medulloblastoma, which is the result of defective base excision repair, and SBS44 in Group4 medulloblastoma, which is also caused by defective DNA mismatch repair. For these signatures, we have not identified any associated consistent genetic alteration enriched across tumors. However, individual tumors harbor non-synonymous somatic mutations in different genes involved in DNA mismatch repair and nucleotide excision repair pathways (e.g. *TP53, POLD1, SMG1, TP73, SWI5*; Supp. Table 6). The possibility that tumors with SBS21, SBS36 and SBS44 activity might have epigenetic alterations affecting known genes in repair pathways and/or mutations in genes that play a role in repair pathways, but for which their function is not yet well characterized cannot be excluded.

Next, we sought to identify any associations between SBS signature activity, chromothripsis and genomic instability. In a cross-cohort analysis, mutations attributed to SBS8 and SBS40 were significantly higher in chromothriptic tumors (Supp. Fig. 3g and Supp. Table 12 for an overview of samples with chromothripsis). Individual cancer type analysis revealed that SBS40 activity in SHH-subtype medulloblastoma was significantly different depending on the presence of chromothripsis in the tumor genome (Supp. Fig. 3g). Genomic instability, quantified here as the total number of structural variants (deletions, duplications, inversions and translocations) identified in a tumor, is highly correlated with SBS2, SBS13 and SBS40 in a cross-cohort analysis (Supp. Fig. 3i). A separate per-cancer type analysis revealed a high correlation of genomic instability and SBS40 activity in SHH and Group4 medulloblastoma, as well as in neuroblastoma (Supp. Fig. 3i). These observations may indicate that mechanisms involved in genomic rearrangements contribute to mutational processes underlying APOBEC signatures and SBS40.

A total of nine small insertion/deletion (ID) signatures were identified from our PedPanCan cohort that matched with the COSMIC v.3 ID signatures (Fig. 2b, Supp. Table 7). As also observed in the PCAWG mutational signature analysis, ID1, ID2, ID5 and ID9 were active across tumors of multiple cancer types (Fig. 2b, Supp. Fig. 2b)^14^. ID1 and ID2, which are the result of DNA damage induced by replication slippage, were present in ∼95% and ∼61% of the tumors in this cohort, respectively (the most prevalent ID signatures in this cohort). Although ID5 and ID9 were present in multiple cancer types, mutations were attributed to these signatures in only a small percentage of tumors (Supp. Table 8, Supp. Table 9). ID8, a signature caused by the potential damage induced by DNA double-strand break repair by non-homologous end joining was present in 6% (n=26) of the whole cohort. The large cohort analysis of tumors as part of PCAWG revealed that ID8 activity was correlated with age at diagnosis, suggesting a clock-like behavior of this signature^14^. However, we did not observe such correlation in our cohort, potentially due to the very low number of tumors with ID8 activity. ID12, a signature with unknown etiology was identified in a small fraction of high-grade glioma and Group3 medulloblastoma. Manual review of aligned reads and variant calls of ID12 assigned indels showed that these mutations were mostly present in low-mappability, repeat-rich regions of the genome, leading to the hypothesis that ID12 might present an artefact signature in this cohort at least.

Next, we sought to identify potential associations of indel signature activity with age, *TP53* mutation status, chromothripsis and genomic instability. Of the 9 ID signatures active in our cohort, we observed that ID1 and ID2 signature activity is highly correlated with age at diagnosis (Supp. Fig. 4a), revealing their clock-like behavior in pediatric cancers as observed in adult cancers^14^. The number of mutations attributed to ID2 and ID9 (of unknown etiology) were significantly different depending on the *TP53* mutation status. *TP53* germline/somatic mutated tumors showed higher activity of ID2 and ID9 compared with wildtype tumors (Supp. Fig 4b). Similarly, a significantly higher number of mutations were attributed to ID2 and ID9 in chromothriptic tumors compared to non-chromothriptic tumors (Supp. Fig. 4c). A similar difference was observed in adult cancers^25^. The total number of structural variants was highly correlated with ID9 signature activity across the cohort. A per-cancer type analysis revealed that ID9 and genomic instability were highly correlated in RMS (Supp. Fig. 4d). In addition, ID3 and ID4 activities were observed in a small fraction of tumors (Supp. Table 8), however, we did not identify any common genomic alterations across these tumors.

Finally, we performed a correlation analysis of SBS and ID signature activities across the cohort to understand if any of the SBS and ID signatures co-occurred. We observed a high correlation amongst clock-like signatures SBS1, SBS5 and ID1, ID2 (Fig. 2c). In addition, ID3, a tobacco smoking related signature was highly correlated with SBS31 and SBS35, both of which are due to DNA damage caused by platinum treatment (Fig. 2b, c).

In summary, we extracted 27 SBS signatures and 9 ID signatures in our pediatric pan-cancer cohort that overlap COSMIC v.3 signatures. The total number of identified signatures is lower than those identified in adult cancers^14^ and a large fraction of mutations (57% and 42% of SBS mutations; 85% and 100% of ID mutations in non-hyper mutated and hyper-mutated samples, respectively) were attributed to clock-like signatures such as SBS1, SBS5, ID1 and ID2. A significant difference in terms of mutations attributed to SBS2, SBS8 and SBS13 as well as ID2 and ID9 was observed to be depending on *TP53* mutation status.

### Homologous recombination repair defect signature activities

The homologous recombination (HR) repair pathway is an error-free mechanism to repair DNA double-strand breaks^26^. Genomic alterations in components of the HR pathway, mainly in the *BRCA1/2* genes, lead to a characteristic pattern of single base substitution mutations and large deletions at microhomology regions, a phenotype frequently termed ‘BRCAness’^27^. Previous studies focusing mainly on breast and ovarian cancers have identified a strong correlation between *BRCA1/2* biallelic pathogenic mutations and activity of signature 3 (COSMIC v.2). However, a significant fraction of these tumors showed signature 3 activity without any identifiable alterations in HR pathway components^5,11,28,14^. Discerning the activity of HR defect COSMIC v.3 signatures SBS3 and ID6 in tumors is important as it has previously been shown to be associated with the therapeutic response to platinum and poly ADP-ribose polymerase (PARP) inhibitor treatment^29,12,30,31,27,28,46^.

Previous analyses of mutational signatures in pediatric cancers, including our own published analysis^2^, have identified a significant proportion of Signature.3 (COSMIC v.2), mostly in tumors without any HR pathway gene defects^2^. However, the current analysis with COSMIC v.3 signatures identified only a small fraction (2.23% of the whole cohort) of tumors with SBS3 signature activity (Fig. 3a,b). This marked difference is most likely the result of previous Signature 3 mutations now being attributed to “flat” signatures (e.g., SBS5 and SBS40) of the updated and refined COSMIC v.3 mutational signatures. In addition, there is a difference in the approach compared to the initial signature analysis, as we assume SBS1 and SBS5 as background signatures and only add SBS3 if it improves the cosine similarity with at least 0.02. However, none of these tumors with SBS3 showed the associated ID6 signature activity, represented by a high fraction of long deletions at microhomology regions (Fig. 2a, Fig. 3b). This lack of ID6 prompted us to test whether our current variant calling pipeline could be penalizing deletions at microhomologies by assigning low confidence, for example, or if pediatric cancers in general have very low numbers of microhomology-associated deletions as an inherent property. In order to test this further, we whole-genome sequenced five tumors from the INFORM registry^32^ with known (likely) pathogenic germline mutations in *BRCA1/2* based on ClinVar. Amongst these, INF_R_1076 had a *BRCA2* pathogenic homozygous mutation and INF_R_025 a *BRCA2* compound heterozygous mutation (both patients also had a clear phenotype, i.e. Fanconi anaemia), while the remaining tumors had heterozygous (likely) pathogenic *BRCA1/2* mutations (Fig. 3c) without any second hit in the tumor. Mutational signature analysis of these five pediatric tumors revealed that only the two tumors with compound heterozygous and homozygous *BRCA2* mutations, i.e. biallelic inactivation, showed SBS3 and ID6 signature activity (Fig. 3c, Supp. Table 10). In addition, we analyzed 22 whole genome sequenced adult tumors from the NCT-MASTER precision oncology program (https://www.nct-heidelberg.de/master) with respect to ID signatures, for which somatic mutations were called with the same DKFZ in-house pipeline. Amongst these, half of the tumors (n=11) had known *BRCA1/2* deficiency and showed clear ID6 activity (Supp. Fig. 5a, Supp. Table 11). These results indicate that our variant calling pipeline does not systematically miss the deletions at microhomology regions and confirm the importance of biallelic *BRCA1/2* inactivation for the presence of predictive HR defect-associated mutational signatures, that is SBS3 and ID6.

**Figure 3:**
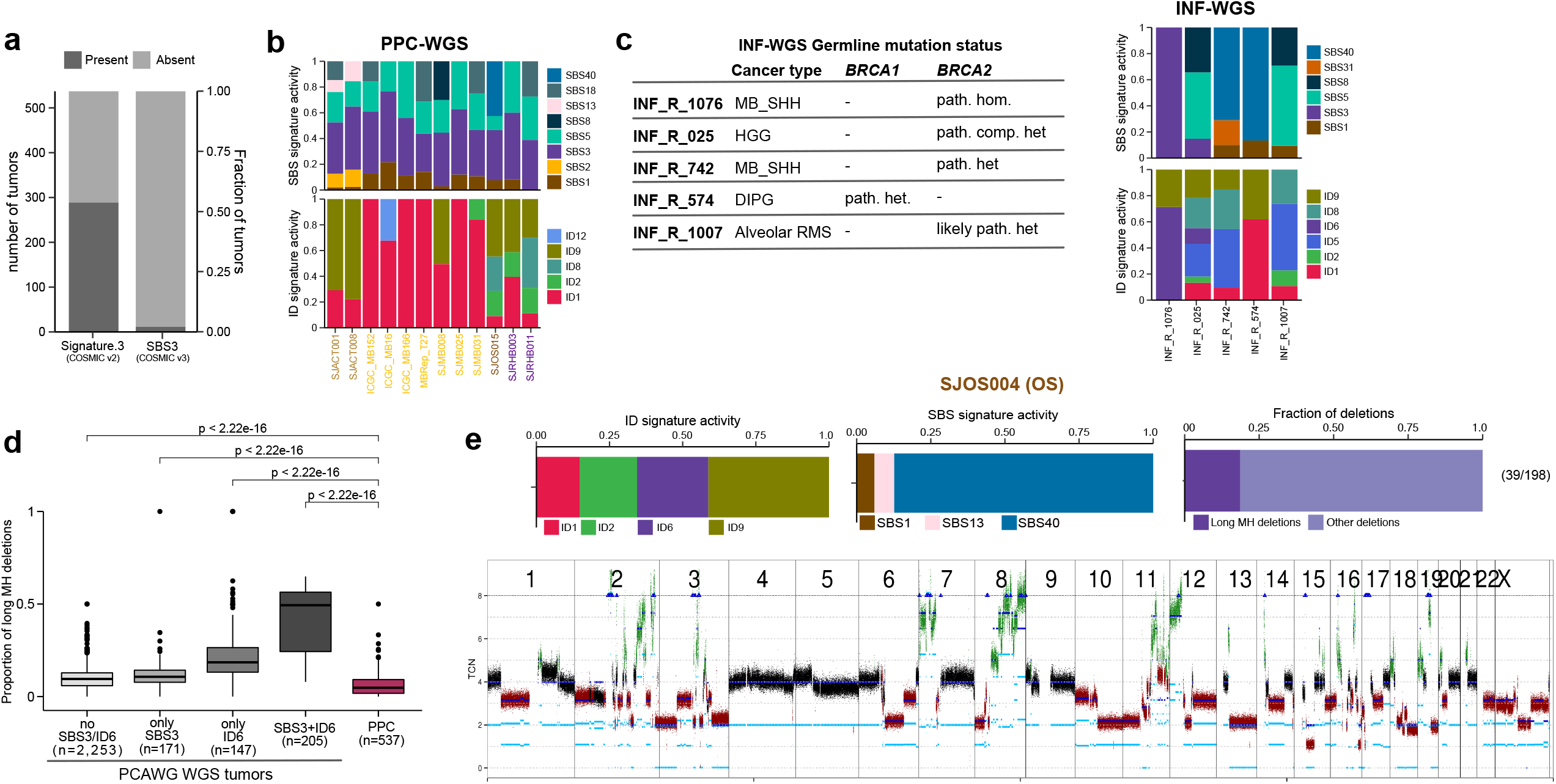
HR defect signatures activity in pediatric cancers. a) Fraction of all PPC-WGS cohort samples that show activity of Signature.3 (COSMIC v.2) and SBS3 (COSMIC v.3). b) Normalized SBS and ID signature activities in tumors with SBS3 signature activity c) BRCA1/2 mutation status of 5 INFORM whole genome sequenced tumors and their SBS and ID signature activities (MB_SHH – Sonic hedgehog subgroup of medulloblastoma; HGG – high-grade glioma; DIPG – diffuse intrinsic pontine glioma; RMS - rhabdomyosarcoma) d) Comparison of microhomology deletion proportion between PCAWG and PPC-WGS cohorts e) SBS, ID signature activities, microhomology deletion proportions and genome wide copy-number profile of osteosarcoma tumor SJOS004.

In the Gröbner *et al*. publication^2^, the initial analysis of this pediatric pan-cancer cohort, six patients were identified with a pathogenic/likely pathogenic *BRCA2* germline variant (Supp. Table 13). Five of these patients were heterozygous in the germline and without evidence for biallelic inactivation in the tumor, while one case was compound heterozygous in the germline for *BRCA2* (SJMB012, 8 year old male with SHH-medulloblastoma). For the latter case, one *BRCA2* germline variant (ENSP00000439902.1:p.Val2407GlufsTer60) is not currently reported in ClinVar and the other one as likely pathogenic (variationID: 421014). We did not identify SBS3 or ID6 activity in any of these six tumors.

Next, we sought to understand whether a proportion of long deletions (>5bp) at microhomologies (MH) are present in pediatric tumors, albeit at very low levels. In order to understand how MH-associated deletions are represented in ID6 positive adult tumors, we divided the PCAWG whole genome tumors (n=2,776) into different categories depending on the presence of SBS3 and ID6. Then we compared the proportion of MH-associated deletions of tumors in these categories with our pediatric cancer cohort. This analysis revealed that pediatric tumors overall have very few MH-associated deletions compared with adult tumors and a significantly lower fraction compared with ID6 positive tumors (Fig. 3d). However, in our cohort, one osteosarcoma tumor (SJOS004) showed ID6 signature without any activity of the SBS3 signature (Fig. 3e). Amongst its SBS signatures, a high fraction of SBS40 mutations were observed. SBS3 and SBS40 are relatively flat signatures and have a similar feature distribution (cosine similarity of 0.88). For this reason, the possibility of mis-assigning SBS3 mutations to SBS40 cannot be excluded. However, no HR pathway mutations were identified in this osteosarcoma tumor.

Amongst the tumors with SBS3 signature activity in our cohort, 58% (7/12) belong to the Group3 subgroup of medulloblastoma (MB). MB tumors are known to follow a linear increase of somatic mutation burden with age^33^. Presumably, mutations in tumors that do not follow such a correlation could be contributed by mutational processes other than the ubiquitous clock-like signatures. In this analysis, we identified only one tumor (SJMB008) with an active SBS3 signature as an outlier of age versus mutation burden correlation, suggesting that most of the other SBS3-positive Group3 MBs follow the typical linear increase in mutation burden with age (Supp. Fig. 5b), and therefore the possibility of mis-assigning mutations from clock-like signatures (especially SBS5 and SBS40) to SBS3 cannot be excluded. Next, we compared expression of HR pathway genes such as *BRCA1/2* and *PALB2* between SBS3 positive and negative tumors in Group 3 MBs, as there could be other mechanisms acting to inactivate the expression of these genes. We observed only a slight difference in *PALB2* expression (Supp. Fig. 5c), but not in the other genes and also no significant difference in promoter DNA methylation of these genes (data not shown).

In summary, we identified only a small percentage (2.23%) of pediatric tumors in our cohort with SBS3 (COSMIC v.3) activity, the so-called HR deficient phenotype, compared to ∼54% of tumors with Signature 3 (COSMIC v.2) activity in our previous analysis of the same cohort^2^. Furthermore, in tumors with SBS3 signature activity, we could not identify any genomic alterations or loss of expression of HR pathway genes. The genome of these SBS3 positive tumors was relatively stable with very few outliers (Supp. Fig. 6). The indel signature ID6, which is characterized by long deletions (>5bp) at microhomology sequences, is a strong predictor of HR deficiency and was not identified in our cohort except for one osteosarcoma tumor. Further analysis of n=5 INFORM and n=22 NCT-MASTER whole genome sequenced tumors suggests the importance of genetic biallelic *BRCA1/2* inactivating events in the generation of HR deficient mutational signatures.

### Signature.P1 similarity to COSMIC v.3 signatures

Previous mutational signature analysis of this cohort based on the COSMIC v.2 reference signatures identified a novel substitution signature, called Signature P1, which featured elevated mutations of C>T in the context of CCC/CCT^2^. Signature P1 was active in the pediatric brain tumors atypical teratoid/rhabdoid tumors (ATRT) and ependymoma. In the current analysis, the Signature P1 profile was compared to all identified COSMIC v.3 SBS signatures and a high similarity was observed with signature SBS31 (Fig. 4a,b). SBS31 is the result of DNA damage caused by platinum treatment. In the present analysis, SBS31 activity was identified in ATRT, ependymoma and ETMR tumors (Fig. 4c). These results suggest that Signature.P1 is not, as previously hypothesized, a pediatric specific mutational signature, but rather a treatment-associated signature identified in a small fraction of tumors that were annotated as treatment naïve. These patients had likely been treated prior to genomic analysis, and at least for one ATRT sample (H049-JVCT; high SBS35 activity, Suppl. Table 2) we could follow up with the sample source which confirmed that this tumor was a recurrence and not a primary tumor.

**Figure 4:**
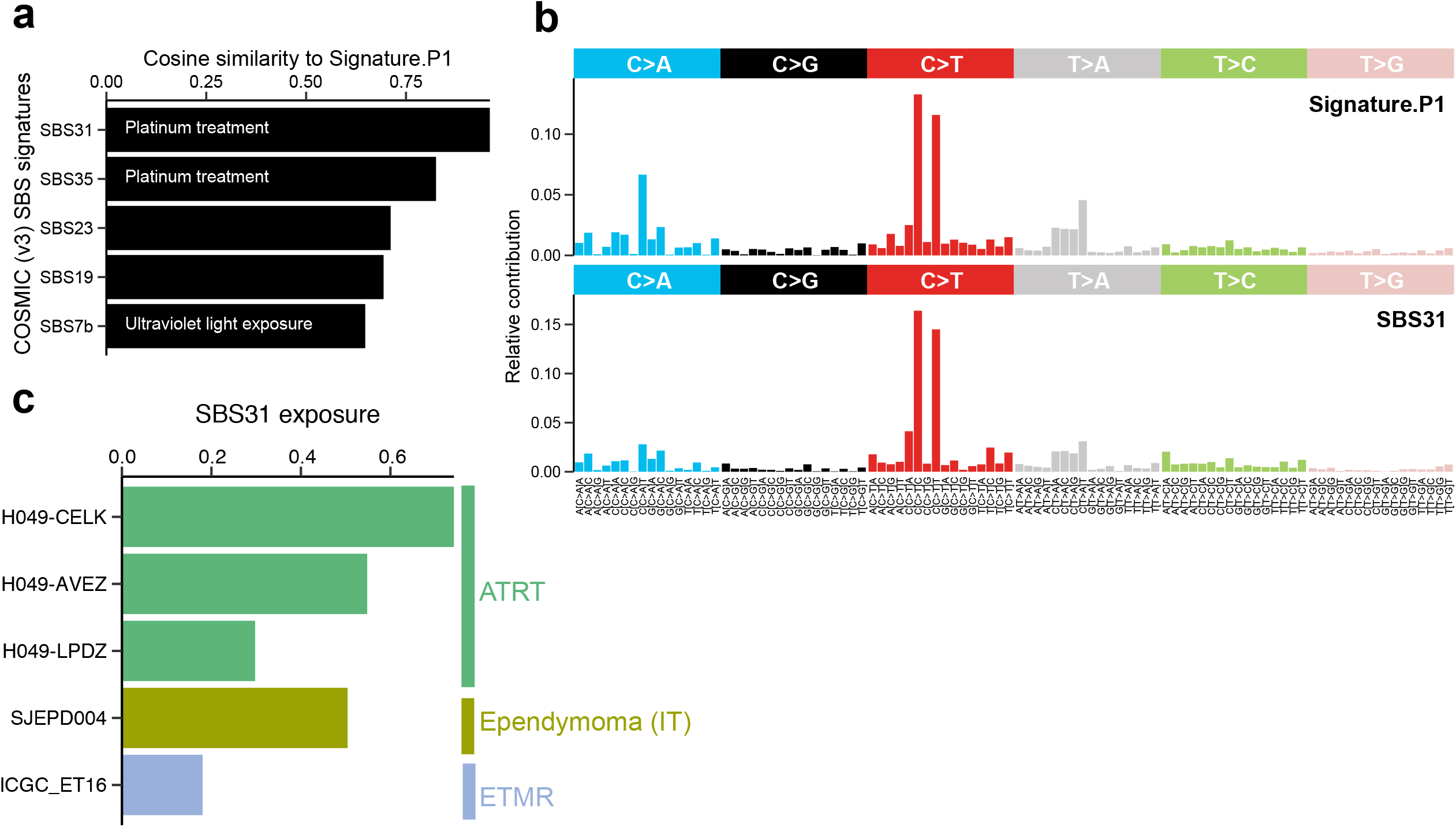
Similarity between Signature.P1 and COSMIC v.3 SBS reference signatures. a) Cosine similarity between novel Signature.P1 from Gröbner, Worst et al., 2018 and COSMIC v.3 SBS signatures. b) 96-context mutational profiles of Signature.P1 and SBS31. c) SBS31 signature activity in the PPC-WGS cohort.

## Discussion

In this study, we re-analyzed 537 whole genome sequenced tumor-normal pairs from 20 different molecularly defined entities of childhood cancers to identify and refine the mutational signatures of the underlying mutational processes. We examined single base substitution (SBS) and small insertion/deletion (ID) in depth and showed that a relatively small number of mutational processes operate in pediatric cancers compared with adult cancers. Amongst the identified 27 SBS and 9 ID signatures, etiologies for more than half of these signatures have been described either by experimental approaches or association analyses in previous studies in adult cancers. In this cohort, a large fraction of SBS (45.4%) and ID (93.2%) mutations across multiple cancer types were attributed to clock-like substitution signatures SBS1 and SBS5,as well as to clock-like indel signatures. Signatures such as SBS8 are relatively frequent across cancer entities, but the etiology remains unknown and recent evidence suggests this signature is due to DNA damage induced by late replication errors^24^. Other signatures without known etiology were identified in a small fraction of tumors from specific cancer types such as pilocytic astrocytoma (namely, SBS12 and SBS23) and Burkitt’s lymphoma (namely, SBS17a, SBS17b). However, we did not identify any genomic alteration common to these tumors.

Previous mutational signature analyses identified Signature 3 (equivalent to the current SBS3) along with other features, with or without discernable alterations in *BRCA1/2* genes, as a potential biomarker for HR deficiency and platinum plus PARPi-based treatment response^34,35^. In pediatric cancers, germline predisposition or somatic alterations in *BRCA1/2* genes are infrequent^2,3^ and only a small fraction (2.23%) of tumors showed SBS3 activity in this analysis, but without the complementing indel signature ID6. Microhomology (MH)-associated deletions, a strong feature of ID6, were significantly more rare in pediatric than in adult cancers. A plausible explanation for this observation could be that the total somatic mutation burden of childhood cancers is very low^2^ and the likelihood of mutations occurring at MH sequences will therefore also be small. Polak *et al*.^11^ identified a signature that is similar to the current SBS3 in breast cancer tumors without alterations in *BRCA1/2* genes, but identified alterations in other HR pathway components (e.g. *PALB2* or *RAD51C*). In this pediatric cancer cohort, we have not identified genomic alterations in other HR pathway genes that could potentially result in the observed SBS3 signature. While there might be other epigenetic mechanisms to inactivate HR pathway genes, in Group3 medulloblastoma (for which complementary omics data is available^33,47^) we have not observed promoter hypermethylation or reduced expression of HR pathway genes. Furthermore, we do not discount the possibility of mis-assigning SBS5 and SBS40 signature mutations to SBS3. In addition, a recent study suggests that SBS3 is likely not as specific as previously believed and that the identification of HR deficiency should rely on multiple orthogonal mutational signatures, not only on SBS3^48^. In order to validate the biomarker efficacy of SBS3 to predict PARP inhibitor treatment response in pediatric cancers, experiments involving PDX models of pediatric tumors are currently ongoing (for example in the “BRCAddict” project, https://www.transcanfp7.eu/index.php/abstract/brcaddict.html). Recent studies focusing on Ewing sarcoma^37^ and ETMR brain tumors ^38^ identified the presence of R-loops, DNA-RNA hybrid structures, to be correlated with PARP inhibitor response. In the future, mutational signature SBS3 combined with other features such as indel signature ID6 and/or R-loops should be investigated in preclinical models to asses PARP inhibitor response and to ultimately define optimized biomarkers.

In summary, although some cancer entities are under-represented in this cohort, we believe this analysis on the mutational signature repertoire in childhood cancers provides a valuable resource for further understanding of tumor biology and aids future research in defining biomarkers of treatment response.

## METHODS

### Whole-genome sequencing data, alignment, and variant calling

All whole genome sequencing data analyzed in this study was collected from Gröbner, Worst et al., 2018^2^. Briefly, FASTQ data was aligned to reference genome GRCh37/hg19 with BWA-MEM (v 0.7.8)^39^. Single base substitutions were called using an updated samtools^40^ based DKFZ in-house pipeline (0.1.19) and indels were called using Platypus (0.8.1.1).

### Somatic Mutation Frequencies

Single base substitution and small insertion/deletion mutation burden was calculated as the total number of mutations identified per megabase of the genome. For whole genome sequencing data, the total number of mutations was divided by 2800 (effective human genome size in megabases that can be assessed by whole-genome sequencing).

### Analysis of mutational signatures using SigProfiler

In order to extract mutational signatures, 96-context SBS and 83-context ID mutational catalogues were prepared using SigProfilerMatrixGenerator (version 1.0.24)^41^ for each of the 20 cancer types. These mutational catalogues were used as input to SigProfilerExtractor (version 1.0.19)^42^ to extract signatures and attribute mutations to each signature in every individual tumor. Briefly, SigProfilerExtractor applies a nonnegative matrix factorization algorithm in multiple iterations (n=100) and for each iteration the software minimizes a generalized Kullback-Leibler divergence constrained for non-negativity. The optimal number of mutational signatures was selected based on highest average stability and lowest average sample cosine distance. After the optimal number of signatures was estimated, attribution of mutations to each signature in each sample involved finding the minimum of the Frobenius norm of a constrained function using a nonlinear convex optimization programming solver based on the interior point algorithm^21^.

### Structural variants and copy-number profiles

Structural variants per tumor were identified as described previously using the DELLY^43^ ICGC pan-cancer analysis workflow (https://github.com/ICGC-TCGA-PanCancer/pcawg_delly_workflow). Copy-numbers were estimated using the ACEseq (allele-specific copy-number estimation from sequencing) too, based on binned tumor-control coverage ratio and B-allele frequencies (BAF)^44^.

## Statistical analyses and code availability

All downstream statistical analyses were performed using the R statistical programming language (version 4.0.3) and all code is available at https://github.com/KiTZ-Heidelberg/Signatures-Manuscript

## Acknowledgements

This project was supported by funding from the ADDRess consortium (Grant 01GM1909E, BMBF). L.B.A. is an Abeloff V scholar and he is personally supported by a Packard Fellowship for Science and Engineering. We acknowledge the DKFZ’s ODCF and GPCF core facilities for supporting the genomic sequencing and data processing.

## Contributions

V.T., S.M.P. and N.J. conceptualized the study. S.M.A.I. and L.B.A. contributed to software development and implementation. Analyses were performed by V.T., N.J., G.W., S.M.A.I and L.B.A. Data visualization was conducted by V.T. Data were collected and curation of data was conducted by V.T., B.C.J., S.N.G., B.H., D.H., S.F., M.B.-J., and D.T.W.J. The original draft was written by V.T. and N.J., and all co-authors reviewed and edited the manuscript.

## Figure Legends

**Supp. Figure 1:**
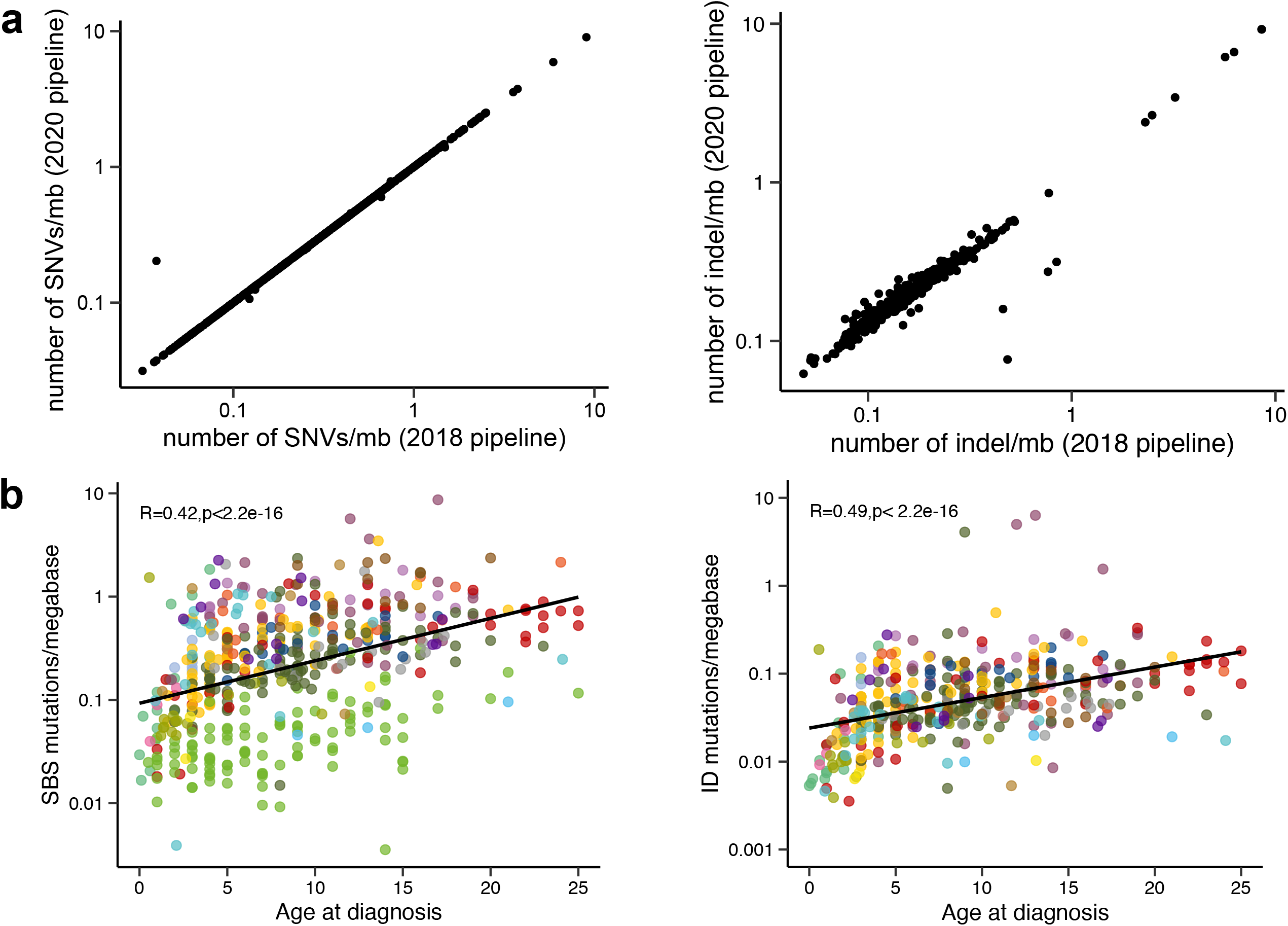
Correlation of SBS and ID mutations with age at tumor diagnosis. a) Comparison of number of mutation calls identified between 2018 (in Gröbner, Worst et al., 2018) and the updated 2020 DKFZ in-house pipeline. b) Correlation of SBS and ID mutations with age across the PPC-WGS cohort.

**Supp. Figure 2:**
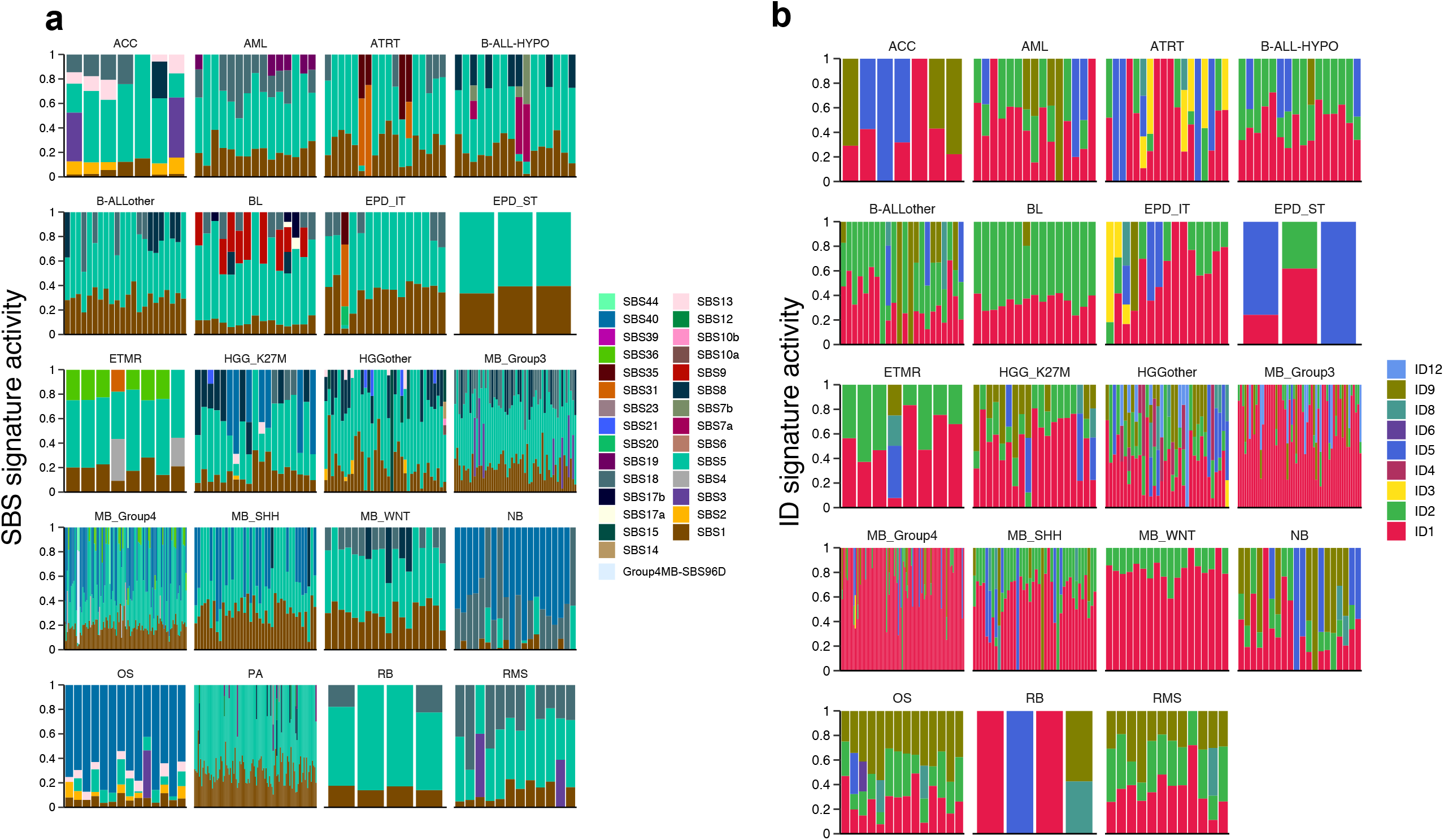
Normalized mutational signature activities across the cohort per sample. a) Normalized SBS signature activities across the cohort b) Normalized ID signature activities

**Supp. Figure 3:**
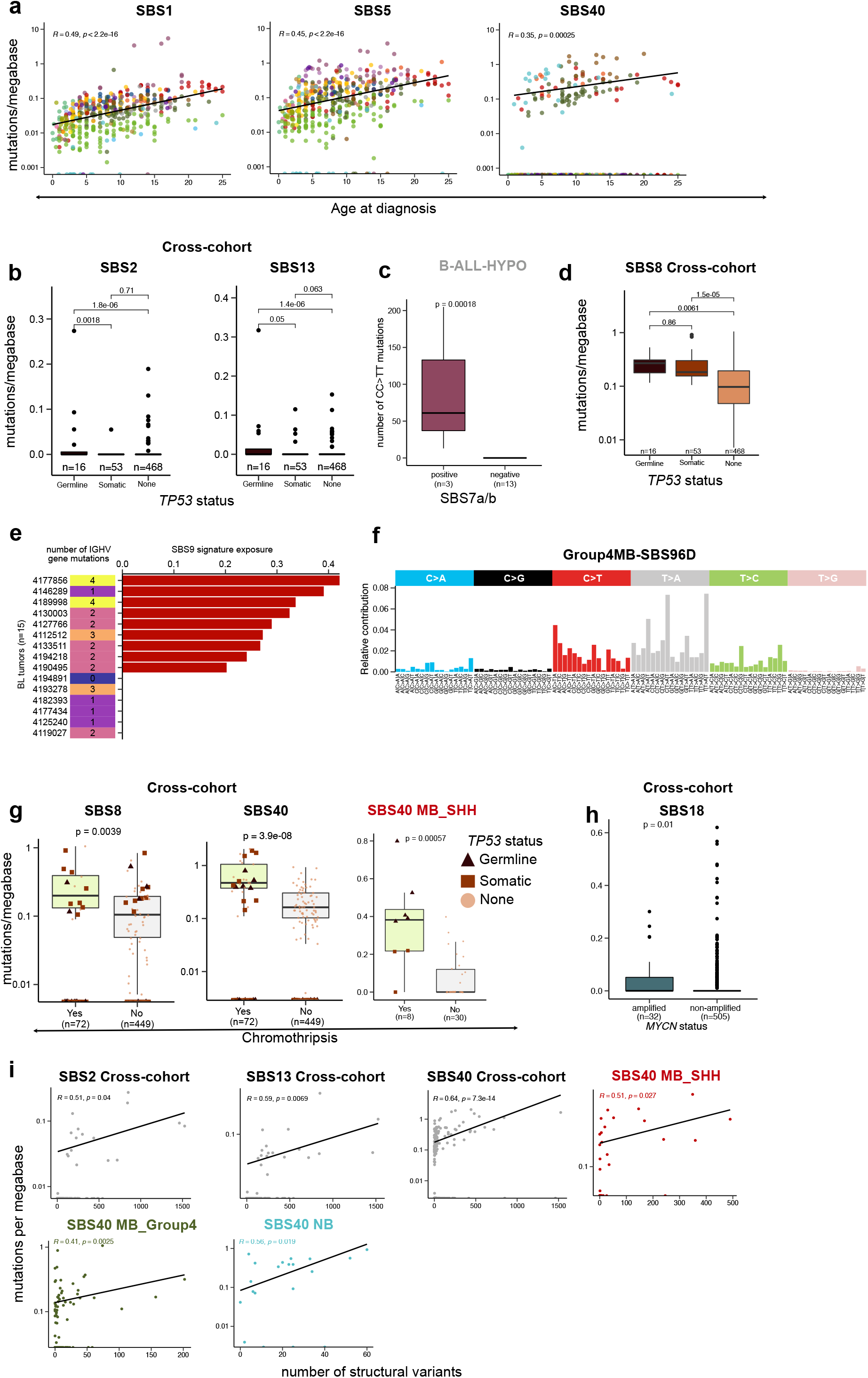
Association of SBS signature activities and genomic alterations. a) Correlation of SBS1, SBS5 and SBS40 signature activities and age at diagnosis. b) Association of *TP53* mutation status and SBS2 and SBS13 signature activities c) Difference in number of CC>TT double base substitutions between SBS7a/b positive and negative tumors of B-ALL-HYPO tumors. d) Association of *TP53* mutation status and SBS8 signature activity e) Number of IGHV mutations and SBS9 signature activities in Burkitt’s lymphoma samples. f) Profile of the novel SBS signature identified in Group4-subgroup of medulloblastoma (Group4MB-SBS96D) g) Association of chromothripsis and SBS8 and SBS40 signature activities. h) Association of MYCN-amplification status and SBS18 signature activity i) Correlation of genomic instability and SBS2, SBS13 and SBS40 signature activities.

**Supp. Figure 4:**
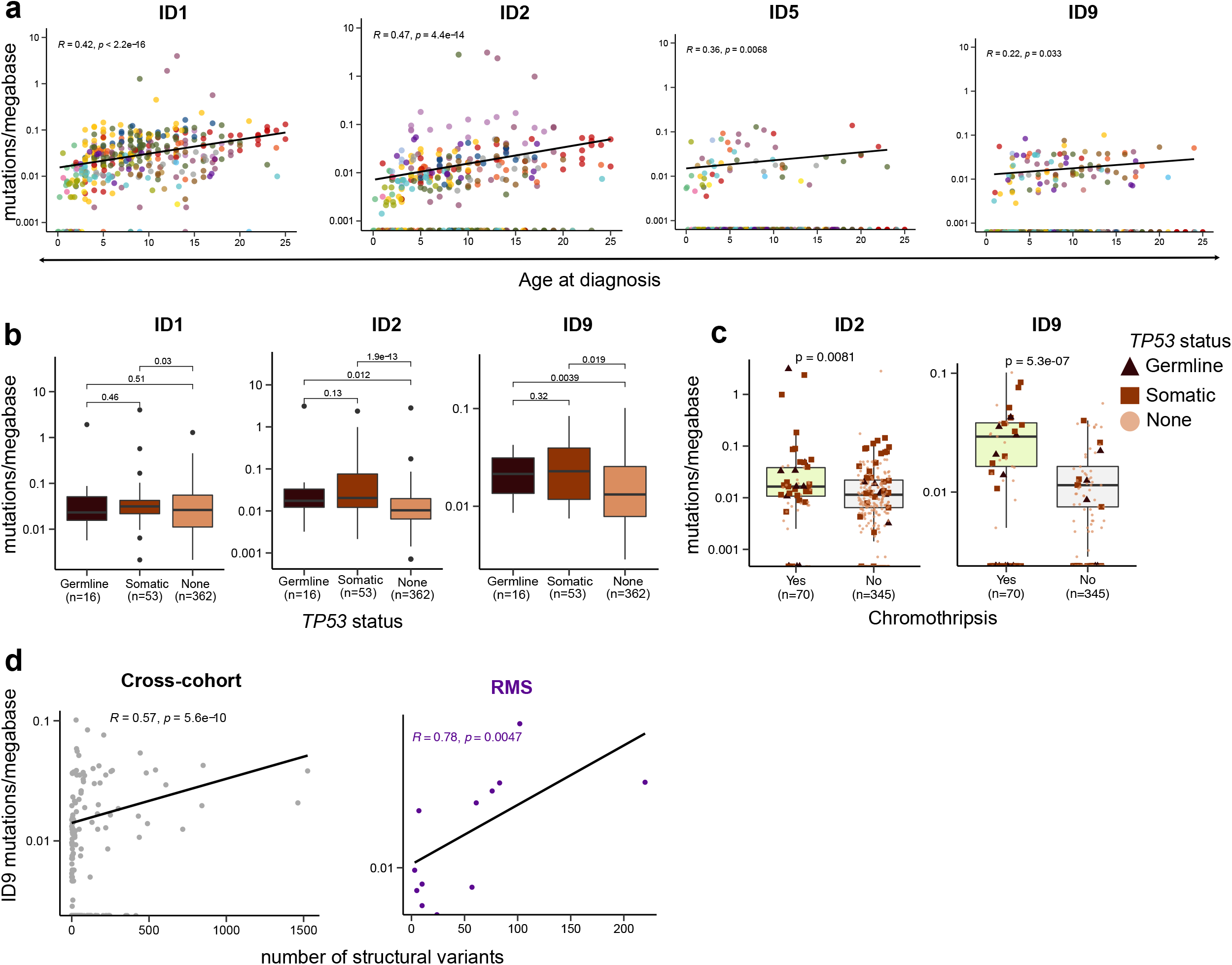
Association of ID signature activities and genomic alterations. a) Correlation of ID1, ID2, ID5 signature activities and age at diagnosis b) Association of TP53 mutation status and ID1, ID2 and ID9 signature activities c) Association of chromothripsis and ID2, ID9 signature activities d) Correlation of genomic instability and ID9 signature activity

**Supp. Figure 5:**
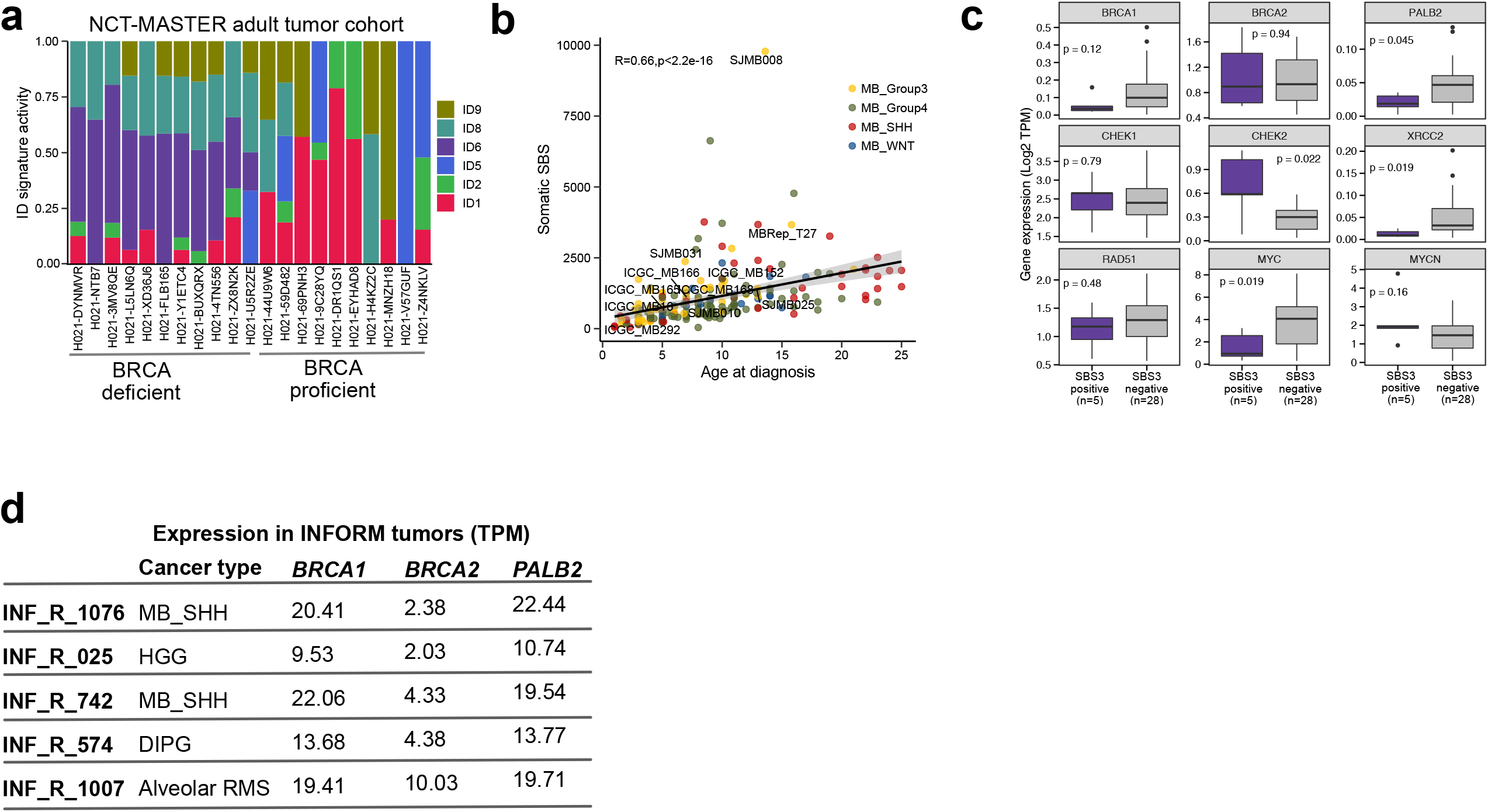
ID signatures in adult cohort and downstream analysis of SBS3 positive tumors. a) ID signature activities in adult tumors of NCT-MASTER b) Correlation of SBS mutation burden and age at diagnosis in medulloblastoma c) HR pathway gene expression difference between SBS3 positive and negative tumors of Group3 subgroup of medulloblastoma d) HR pathway gene expression in INFORM whole genome sequenced tumors

**Supp. Figure 6:**
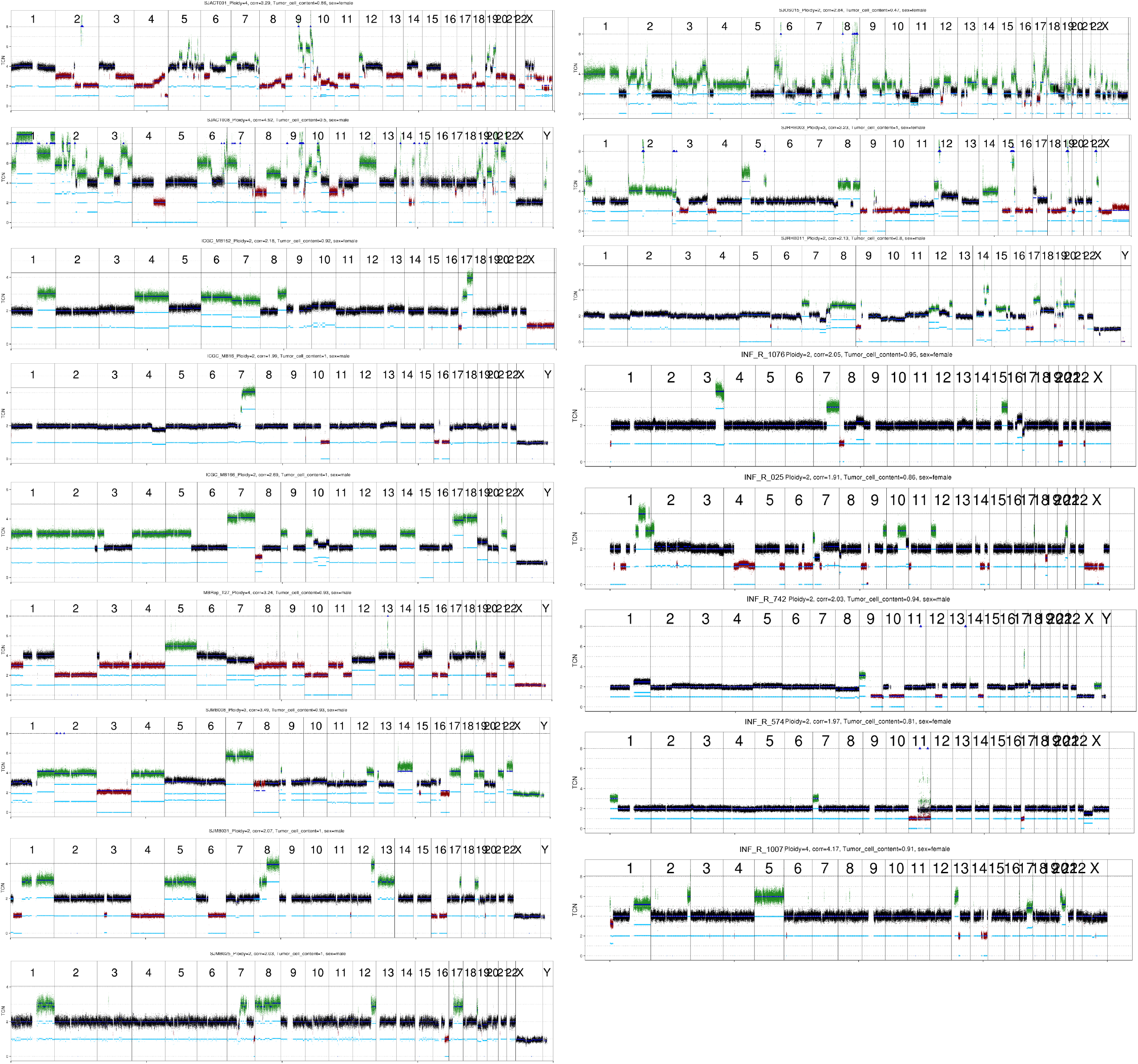
Genome-wide copy-number profiles of SBS3 positive tumors from the PedPanCan cohort and INFORM tumors (n=5) TCN = Tumor Copy Number. Red indicates loss of genomic material, green indicates gains. Plots were generated using the ACEseq tool.

## Notes

### Competing Interest Statement

The authors have declared no competing interest.

## REFERENCES

1. Pui, C., Gajjar, A. et al. Challenging issues in pediatric oncology. Nat. Rev. Clin. Oncol 540, 540–549 (2011).

2. Gröbner, S., Worst, B., Weischenfeldt, J. et al. The landscape of genomic alterations across childhood cancers. Nature 555, 321–327 (2018)

3. Ma, X. et al. Pan-cancer genome and transcriptome analyses of 1,699 paediatric leukaemias and solid tumours. Nature 555, 371–376 (2018).

4. Stratton, M. R., Campbell, P. J. & Futreal, & P. A. The cancer genome. Nature 458, (2009).

5. Alexandrov, L., Nik-Zainal, S., Wedge, D. et al. Signatures of mutational processes in human cancer. Nature 500, 415–421 (2013)

6. Nik-Zainal, S. et al. Mutational processes molding the genomes of 21 breast cancers. Cell 149, 979–993 (2012).

7. Alexandrov, L., Jones, P., Wedge, D. et al. Clock-like mutational processes in human somatic cells. Nat Genet 47, 1402–1407 (2015).

8. Schulze, K., Imbeaud, S., Letouzé, E. et al. Exome sequencing of hepatocellular carcinomas identifies new mutational signatures and potential therapeutic targets. Nat Genet 47, 505–511 (2015).

9. Nik-Zainal, S. et al. Landscape of somatic mutations in 560 breast cancer whole-genome sequences. Nature 534, 47–54 (2016).

10. Petljak, M. & Alexandrov, L. B. Understanding mutagenesis through delineation of mutational signatures in human cancer. Carcinogenesis 37, 531–540 (2016).

11. Polak, P. et al. A mutational signature reveals alterations underlying deficient homologous recombination repair in breast cancer. Nat. Genet. 49, 21 (2017).

12. Keung, M., Wu, Y. & Vadgama, J. PARP Inhibitors as a Therapeutic Agent for Homologous Recombination Deficiency in Breast Cancers. J. Clin. Med. 8, 435 (2019).

13. Campbell, P. J. et al. Pan-cancer analysis of whole genomes. Nature 578, 82–93 (2020).

14. Alexandrov, L. B. et al. The repertoire of mutational signatures in human cancer. Nature 578, 94–101 (2020).

15. Johann, P. D. et al. Atypical Teratoid/Rhabdoid Tumors Are Comprised of Three Epigenetic Subgroups with Distinct Enhancer Landscapes. Cancer Cell 29, 379–393 (2016).

16. Northcott, P., Buchhalter, I., Morrissy, A. et al. The whole-genome landscape of medulloblastoma subtypes. Nature 547, 311–317 (2017)

17. Kovac, M. et al. Exome sequencing of osteosarcoma reveals mutation signatures reminiscent of BRCA deficiency. Nat. Commun. 6, 1–9 (2015).

18. Kool, M. et al. Genome sequencing of SHH medulloblastoma predicts genotype-related response to smoothened inhibition. Cancer Cell 25, 393–405 (2014).

19. Kunz, J. B. et al. Pediatric T-cell lymphoblastic leukemia evolves into relapse by clonal selection, acquisition of mutations and promoter hypomethylation. Haematologica 100, 1442–1450 (2015).

20. Jones, D. T. W. et al. Recurrent somatic alterations of FGFR1 and NTRK2 in pilocytic astrocytoma. Nat. Genet. 45, 927–932 (2013).

21. Alexandrov, L. B., Nik-Zainal, S., Wedge, D. C., Campbell, P. J. & Stratton, M. R. Deciphering Signatures of Mutational Processes Operative in Human Cancer. Cell Rep. 3, 246–259 (2013).

22. Periyasamy, M. et al. p53 controls expression of the DNA deaminase APOBEC3B to limit its potential mutagenic activity in cancer cells. Nucleic Acids Res. 45, 11056–11069 (2017).

23. Wang, S., Jia, M., He, Z. & Liu, X.-S. APOBEC3B and APOBEC mutational signature as potential predictive markers for immunotherapy response in non-small cell lung cancer. Oncogene 37, 3924–3936 (2018).

24. Singh, V.K., Rastogi, A., Hu, X. et al. Mutational signature SBS8 predominantly arises due to late replication errors in cancer. Commun Biol 3, 421 (2020).

25. Voronina, N., Wong, J.K.L., Hübschmann, D. et al. The landscape of chromothripsis across adult cancer types. Nat Commun 11, 2320 (2020).

26. Jasin, M. & Rothstein, R. Repair of Strand Breaks by Homologous Recombination. (2013) doi:10.1101/cshperspect.a012740

27. Byrum, A. K., Vindigni, A. & Mosammaparast, N. Defining and Modulating ‘BRCAness’. Trends in Cell Biology 29, 740–751 (2019).

28. Davies, H. et al. HRDetect is a predictor of BRCA1 and BRCA2 deficiency based on mutational signatures. Nat. Med. 23, 517–525 (2017).

29. Póti, Á. et al. Correlation of homologous recombination deficiency induced mutational signatures with sensitivity to PARP inhibitors and cytotoxic agents. Genome Biol. 20, 240 (2019).

30. Chen, A. PARP inhibitors: its role in treatment of cancer. Chinese journal of cancer 30, 463–471 (2011).

31. Tutt, A. et al. Oral poly(ADP-ribose) polymerase inhibitor olaparib in patients with BRCA1 or BRCA2 mutations and advanced breast cancer: A proof-of-concept trial. Lancet 376, 235–244 (2010).

32. Worst, B. C. et al. Next-generation personalised medicine for high-risk paediatric cancer patients – The INFORM pilot study. Eur. J. Cancer 65, 91–101 (2016).

33. Jones, D. T. W. et al. Dissecting the genomic complexity underlying medulloblastoma. Nature 488, 100–105 (2012).

34. Ma, J., Setton, J., Lee, N. Y., Riaz, N. & Powell, S. N. The therapeutic significance of mutational signatures from DNA repair deficiency in cancer. Nature Communications 9, (2018).

35. Zhao, E. Y. et al. Personalized Medicine and Imaging Homologous Recombination Deficiency and Platinum-Based Therapy Outcomes in Advanced Breast Cancer. Clin Cancer Res 23, (2017).

36. Machado, H. et al., Genome-wide mutational signatures of immunological diversification in normal lymphocytes. (2021) bioRxiv 2021.04.29.441939; doi: https://doi.org/10.1101/2021.04.29.441939

37. Gorthi, A. et al. EWS-FLI1 increases transcription to cause R-Loops and block BRCA1 repair in Ewing sarcoma. Nature 555, 387–391 (2018).

38. Lambo, S. et al. The molecular landscape of ETMR at diagnosis and relapse. Naure 576, 576(7786):274-280 (2019).

39. Li, H. & Durbin, R. Fast and accurate short read alignment with Burrows-Wheeler transform. Bioinformatics 25, 1754–1760 (2009).

40. Li, H. et al. The Sequence Alignment/Map format and SAMtools. Bioinforma. Appl. NOTE 25, 2078–2079 (2009).

41. Bergstrom, E. N. et al. SigProfilerMatrixGenerator: A tool for visualizing and exploring patterns of small mutational events. BMC Genomics 20, 685 (2019).

42. Ashiqul Islam, S. M. et al. Uncovering novel mutational signatures by de novo extraction with. (2021) doi:10.1101/2020.12.13.422570, BioRxiv

43. Rausch, T. et al. DELLY: structural variant discovery by integrated paired-end and split-read analysis. Bioinformatics 28, 333–339 (2012).

44. Kleinheinz, K. et al. ACEseq-allele specific copy number estimation from whole genome sequencing. (2017) doi:10.1101/210807, BioRxiv

45. Li, B. et al., Therapy-induced mutations drive the genomic landscape of relapsed acute lymphoblastic leukemia. Blood. 2020; 135(1): 41–55.

46. Gröschel, S., Hübschmann, D., Raimondi, F. et al. Defective homologous recombination DNA repair as therapeutic target in advanced chordoma. Nat Commun 10, 1635 (2019).

47. Hovestadt, V., Jones, D., Picelli, S. et al. Decoding the regulatory landscape of medulloblastoma using DNA methylation sequencing. Nature 510, 537–541 (2014).

48. Degasperi, A., Amarante, T.D., Czarnecki, J. et al. A practical framework and online tool for mutational signature analyses show intertissue variation and driver dependencies. Nat Cancer 1, 249–263 (2020).

